# BORIS/CTCFL promotes a switch from a proliferative towards an invasive phenotype in melanoma cells

**DOI:** 10.1101/560474

**Authors:** Sanne Marlijn Janssen, Roy Moscona, Mounib Elchebly, Andreas Ioannis Papadakis, Margaret Redpath, Hangjun Wang, Eitan Rubin, Léon Cornelis van Kempen, Alan Spatz

## Abstract

Melanoma is among the most aggressive cancers due to its tendency to metastasize early. Phenotype switching between a proliferative and an invasive state has been suggested as a critical process for metastasis, though the mechanisms that regulate state transitions are complex and remain poorly understood. Brother of Regulator of Imprinted Sites (BORIS), also known as CCCTC binding factor-Like (CTCFL), is a transcriptional modulator that becomes aberrantly expressed in melanoma. Yet, the role of BORIS in melanoma remains elusive. Here, we show that BORIS is involved in melanoma phenotype switching. Genetic modification of BORIS expression in melanoma cells combined with whole transcriptome analysis indicated that BORIS expression contributes to an invasion-associated transcriptome. In line with these findings, inducible BORIS overexpression in melanoma cells reduced proliferation and increased migration and invasion, demonstrating that the transcriptional switch is accompanied by a phenotypic switch. Mechanistically, we reveal that BORIS binds near the promoter of transforming growth factor-beta 1 (*TFGB1*), a well-recognized factor involved in the transition towards an invasive state, which coincided with increased expression of *TGFB1*. Overall, our study indicates a pro-invasive role for BORIS in melanoma via transcriptional reprogramming.

## Introduction

Primary cutaneous melanoma is one of the most aggressive cancers due to its ability to rapidly disseminate and develop distant metastases. The progression of melanoma from a primary to metastatic tumor requires melanoma cells to gain invasive abilities, allowing them to migrate and invade into the lymphovascular system and colonize distant organs. During melanoma progression, cells undergo a reversible transition from a proliferative to an invasive state, a process known as “phenotype switching”, which closely resembles epithelial-mesenchymal transition (EMT) (1, 2). Gene expression profiling of melanoma cell lines and tumor samples has revealed the different transcriptional states that underlie the proliferative and invasive phenotypes of melanoma cells (3–7). While high expression of the transcription factor MITF is linked to the proliferative state, TGF-beta was the first factor associated with expression of invasion-associated genes (3). Currently, various other transcriptional regulators have been implicated in the switch towards an invasive state, including BRN2, AP-1, TEAD, NFATC2, and NFIB (4, 8–11). Furthermore, integration of transcriptomic and epigenomic data from melanoma cells revealed widespread differences in the chromatin landscape between the proliferative and invasive cellular states. More specifically, invasive samples demonstrated active histone modifications and open chromatin in the promoter region of invasion-associated genes, while these regions were marked by closed and transcriptionally repressed chromatin in proliferative cells, indicating that the chromatin landscape plays an important role in phenotype switching (4). While these findings have greatly contributed to our understanding of phenotype switching, and therefore to the mechanisms that underlie melanoma metastasis, new insights may be derived from the identification of factors that regulate the transcriptional landscape either directly as a transcriptional regulator or indirectly by changing the chromatin landscape.

Brother of Regulator of Imprinted Sites (BORIS) is a DNA binding protein with high similarity to CCCTC binding factor (CTCF) (12), a multifunctional transcription factor that plays an important role in chromatin organization (13). In contrast to the tumor suppressive functions reported for CTCF (14, 15), multiple studies have indicated an oncogenic role for BORIS (16–20). Like CTCF, BORIS plays a role in transcriptional regulation (17, 18, 21, 22). The mechanisms through which BORIS can alter transcription rely on BORIS’ ability to bind the DNA at specific binding motifs. These motifs are often found near promoter regions where BORIS-DNA binding is enriched when the chromatin harbors activating histone marks and is in an accessible state (23–25). At DNA binding sites BORIS can compete with DNA-bound CTCF (25), form a BORIS-CTCF heterodimer, or occupy a so-called BORIS-only binding site (24). Via these mechanisms BORIS can alter transcription by acting as a transcriptional activator (17, 18, 22, 26), recruiting a transcriptional activator (27), altering DNA methylation (17, 28, 29) or histone modifications (17, 21, 22, 28), or recruiting chromatin modifying proteins (30, 31). In addition, it is believed that BORIS impacts the chromatin landscape by interfering with CTCF-mediated chromatin loops (24, 32).

BORIS expression is normally restricted to the testis and becomes aberrantly expressed in different types of cancer, hence the designation of BORIS as a cancer testis antigen (12). In cancer cells, BORIS expression is frequently facilitated via hypomethylation of the *BORIS* promoter (33, 34). Furthermore, the *BORIS* gene is located at the 20q13 locus, within a chromosomal region that is often amplified in cancer (35). In melanoma, *BORIS* expression was first described by Vatolin *et al*. who detected *BORIS* expression in 9 out of 10 melanoma cell lines (29)). The most elaborate analysis of *BORIS* expression in melanoma was performed by Kholmanskik *et al.* who reported *BORIS* expression in 59% of melanoma cell lines, in 16% (4 out of 25) of primary melanomas and in 34% (13 out of 38) of melanoma metastases, with *BORIS* reaching similar expression levels as observed in the testis (36). Importantly, no *BORIS* expression was observed in normal human skin tissue (36). Although these observations suggest that BORIS can play a role in melanoma progression, little is known about the role of BORIS in melanoma development and progression.

In this study, we sought to determine if BORIS plays a role in melanoma progression through its function as transcriptional modulator. Using a doxycycline (dox)-inducible expression system in melanoma cells we found that BORIS expression led to large-scale changes in gene expression as determined by RNA-sequencing (RNA-seq). While the upregulated genes were enriched among invasion-related processes and gene signatures, the downregulated genes were enriched among proliferation-related processes and gene signatures, suggesting that BORIS plays an important role in the transcriptional switch from a proliferative towards an invasive state. Accordingly, we observed that BORIS overexpression reduced proliferation and increased the migratory and invasive abilities of melanoma cells. Interestingly, we found that BORIS binds in the vicinity of the *TGFB1* promoter, which co-occurs with increased expression of *TGFB1* as well as TGF-beta family members and TGF-beta target genes. Overall, these findings identify BORIS as a mediator of transcriptional reprogramming in melanoma cells, resulting in a switch towards an invasive phenotype.

## Results

### Increased *BORIS* expression in melanoma

To corroborate previous findings that demonstrated *BORIS* expression in human melanoma tumors, but not in normal human skin (36), we downloaded and analyzed *BORIS* expression data for normal testis and skin, and melanoma samples obtained from the Genotype-Tissue Expression (GTEx) and The Cancer Genome Atlas (TCGA) melanoma dataset (SKCM), respectively. *BORIS* expression was detected (log_2_(norm_count+1)>0) in all testis samples, 38.1% of normal skin samples (212 out of 556) and 68.8% of melanoma samples (322 out of 468). We found significantly higher *BORIS* expression in melanoma tumors and normal testis compared to normal skin samples (**Figure S1A**). In addition, the TCGA SKMC dataset revealed significantly higher *BORIS* expression among metastatic melanoma samples compared to primary melanoma samples (**Figure S1B**). Furthermore, in our panel of melanoma and non-malignant congenital nevi cell lines we observed *BORIS* mRNA in almost all of the melanoma cell lines (9 out of 10), but not in the non-malignant congenital nevi cell lines (**Figure S1C**). Collectively, these data confirm that BORIS is overexpressed in melanoma cell lines and tumor samples compared to non-malignant cell lines and normal skin samples, suggesting that BORIS may play an important role in melanoma development and metastasis.

### BORIS expression results in reduced proliferation and increased apoptosis

To gain insight into the role of BORIS in melanoma, we first established a dox-inducible model of BORIS expression in the MM057 melanoma cell line. These cells were used, since they express low *BORIS* mRNA compared to other melanoma cell lines (**Figure S1C)** and have been well characterized by our laboratory. The expression of BORIS in the presence of dox was confirmed by immunoblot (**Figure S2A**). A striking reduction in cell number was observed during cell culture of BORIS-expressing cells [BORIS with dox (BORpos)] compared to the control cells [empty vector without dox (EVneg) and with dox (EVpos), and BORIS without dox (BORneg] (**Figure S2B**). We used this phenotype to optimize the level (dox concentration) and duration of BORIS expression to further characterize the effect of BORIS expression in melanoma cells. To this end, BORIS expression was induced with increasing concentration of dox for either three days, five days or seven days. Expression of BORIS at each time point was confirmed using immunoblot, which demonstrated increasing BORIS expression level with an increasing concentration of dox (**Figure S2C**). Expression of BORIS for three days did not significantly reduce cell proliferation compared to untreated cells, except in the presence of the highest concentration of dox (**Figure 1A**). After five and seven days of BORIS expression, we observed a significant reduction in proliferation even at low dox concentrations (**Figure 1B and C**). Of note, at these concentrations we observed no effect of dox on the proliferation of the empty vector control cells, indicating that doxycycline itself does not lead to reduced proliferation as a result of cell toxicity. Based on these results, we decided to further study the role of BORIS in melanoma cells by inducing BORIS expression with 50-100ng/ml dox for five days of cell culture.

**Figure 1.**
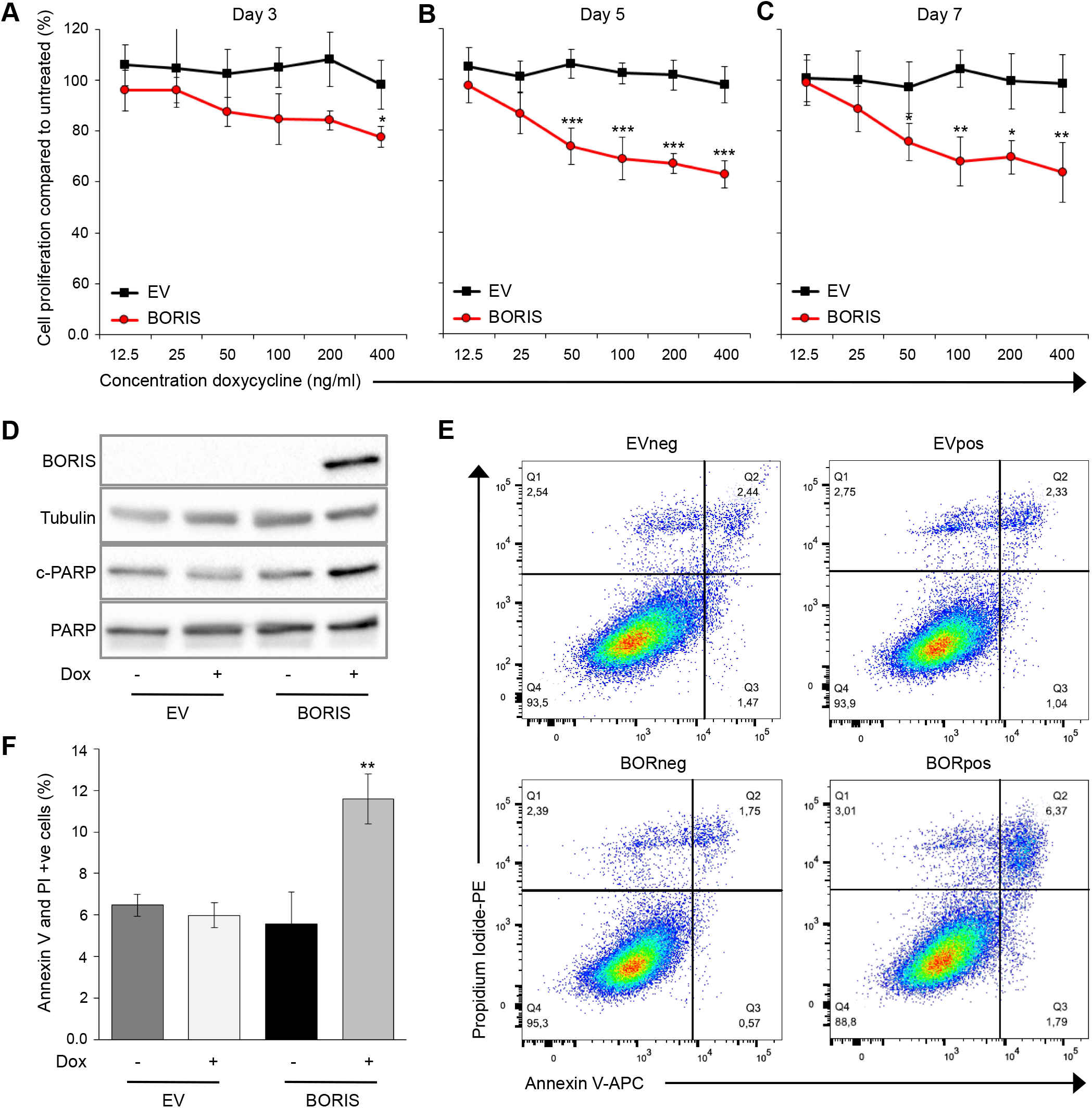
BORIS expression in melanoma cells results in reduced proliferation and increased apoptosis. **(A-C)** Cell proliferation upon expression of BORIS protein in the MM057 cell line with the indicated concentration of dox for 3, 5 or 7 days of cell culture. **(D-F)** BORIS expression was induced in MM057 cells with 50ng/ml dox for 5 days. **(D)** Whole cell lysate was used for immunoblotting with anti-BORIS, anti-cleaved PARP (c-PARP) and anti-PARP antibodies. Anti-Tubulin was used as a loading control. **(E)** Bar graph representing the mean percentage of Annexin V and PI positive cells as determined by flow cytometry. **(F)** Representative image of the percentage of apoptotic and necrotic cells as determined by flow cytometry analysis of Annexin V and PI staining. Error bars represent the standard error of mean. * indicates significantly different from **A-** untreated, **E-** the control (\**P*<0.05. \*\**P*<0.01, \*\*\**P*<0.001). Dox: doxycycline.

As it was previously demonstrated that altered BORIS expression has an effect on apoptosis in breast cancer and colon cancer cell lines (16, 37), we wanted to address whether the reduction in proliferation that we observed is due to BORIS-induced apoptosis. To this end, we induced BORIS expression with 50ng/ml dox and investigated the effect of BORIS expression on cell death. Compared to control cells, BORpos cells demonstrated an increase in expression of cleaved PARP, a marker for cell death (**Figure 1D**). In addition, we observed an increase in the number of early apoptotic, apoptotic and necrotic cells following BORIS expression compared to controls (∼6%) (**Figure 1E and F**), confirming that low level BORIS expression in melanoma cells leads to a slight, though significant, increase in apoptosis. Taken together, our results show that expression of BORIS in melanoma cells leads to decreased proliferative activity (∼25%), which is in part due to increased apoptosis (∼6%).

### BORIS expression induces large-scale differential gene expression

Given that BORIS plays a role in transcriptional regulation (17, 18, 21, 22), we next wanted to investigate the effect of BORIS expression on the transcriptome of melanoma cells. To this end, we induced BORIS expression for five days with 50ng/ml dox in the MM057 cell line and performed RNA-seq (three biological replicates). For each independent RNA-seq sample, BORIS expression was confirmed at the mRNA and protein level in the BORpos cells compared to control cells (**Figure 2A**). Since the RNA-seq data demonstrated a strong correlation between the biological replicates (r≥0.97), all of the RNA-seq data was included for downstream analyses.

**Figure 2.**
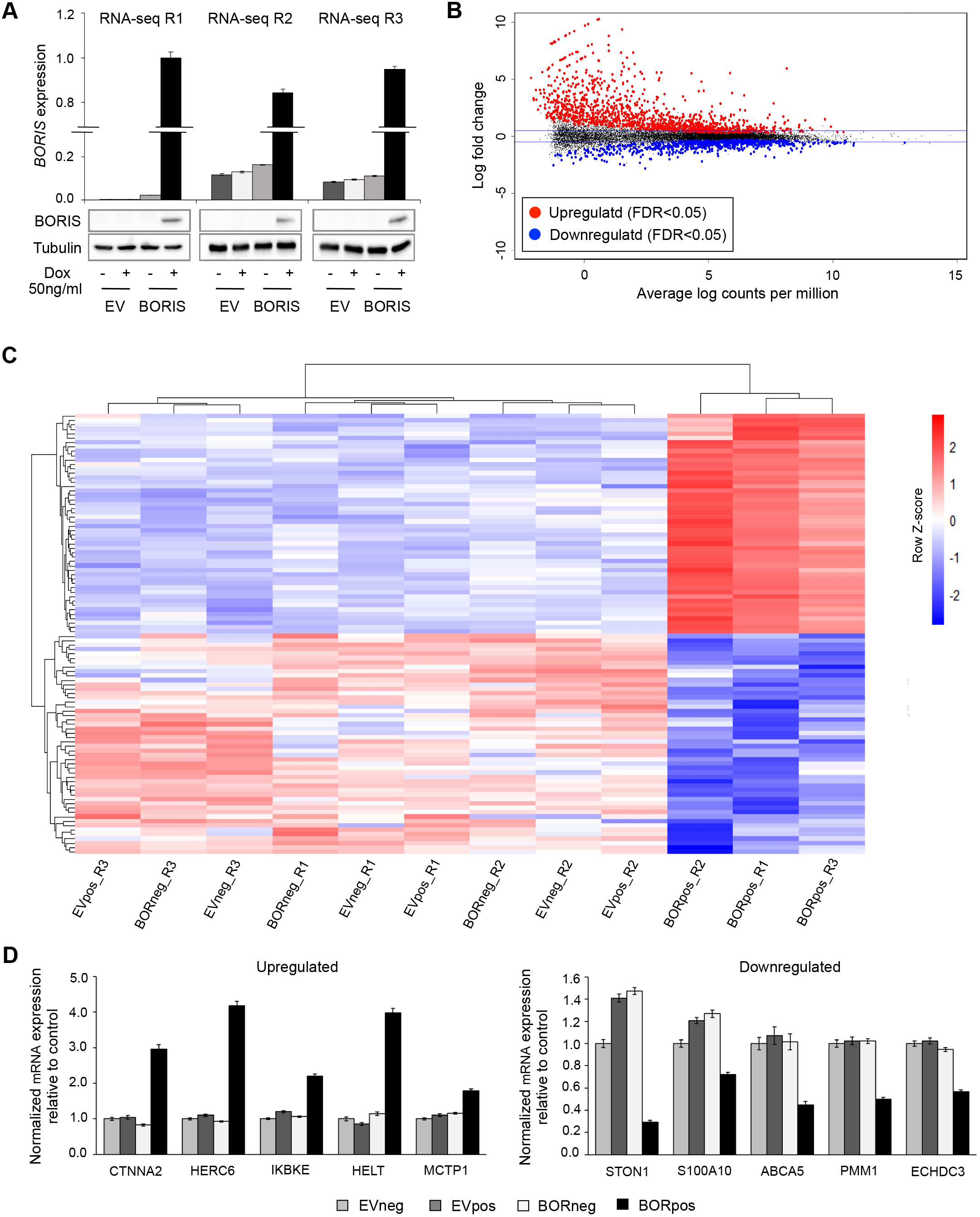
BORIS expression in melanoma cells leads to large-scale differential gene expression. (**A, D**) BORIS expression was induced in the MM057 cell line using 50ng/ml dox for 5 days. **(A)** Expression of BORIS mRNA and protein as determined by qPCR and immunoblot. Anti-Tubulin was used as a loading control for immunoblot. **(B)** MA plot representing the RNA-seq data of all BORneg versus BORpos samples. The blue horizontal lines mark log_2_FC=0.5. **(C)** Heat map representing the log_2_CPM for the top 100 up- and down-regulated genes across all RNA-seq samples. **(D)** qPCR for upregulated and downregulated genes. For each qPCR experiment, a technical triplicate was performed for two biological replicates. Expression was normalized to *HPRT1* and *TBP*. Error bars represent the standard error of the mean. R: replicate, dox: doxycycline, FDR: false discovery rate.

Unsupervised analysis of gene expression demonstrated clustering of the samples into two groups, one representing the three controls (EVneg, EVpos, and BORneg) and the other representing BORIS expressing cells (BORpos) (**Figure S3**). Differential gene expression analysis between BORneg and BORpos cells identified 2045 differentially expressed genes (DEGs) with a false discovery rate (FDR)<0.05 and a |log_2_ fold-change (log_2_FC)|>0.5. From these DEGs, 1308 genes were significantly upregulated and 737 significantly downregulated following BORIS expression (**Figure 2B and Table S2**). Analysis of the log2 counts per million (CPM) for the top 100 up- and down-regulated genes demonstrated robust differential expression, as well as clustering of the three control samples versus BORpos samples (**Figure 2C**). To validate the RNA-seq dataset, BORIS expression was induced in the MM057 cells and qPCR was performed for genes distributed throughout the lists of upregulated and downregulated DEGs. We observed significant differential expression of the selected genes following BORIS expression compared to the controls (**Figure 2D**). Together, these data demonstrate that ectopic BORIS expression in the MM057 melanoma cell line results in large-scale differential gene expression.

It is important to mention that the RNA-seq revealed no effect of increased BORIS levels on the expression of its ubiquitously expressed paralogue *CTCF* (log_2_FC=−0.007, FDR=0.996). In addition, induction of BORIS expression did not change CTCF protein level (**Figure S4A**). Furthermore, we observed no correlation between the expression of *BORIS* and *CTCF* in the TCGA melanoma dataset (R=0.002, *P*=0.96, **Figure S4B**). These data indicate that in neither melanoma cell lines nor patient samples, BORIS expression level has an effect on the expression of CTCF.

### BORIS expression contributes to a switch from a proliferative to invasive transcriptional state

Next, we used the RNA-seq dataset for gene set enrichment analysis (GSEA) (38) to identify biological processes related to melanoma development and progression that BORIS might be involved in. In agreement with our previous observation regarding proliferation (**Figure 1B and C**), we observed a strong negative enrichment (FDR<0.01) for proliferation-related processes like cell cycle/division, telomere maintenance, and DNA replication/repair (**Figure 3A**). Interestingly, we found a strong positive enrichment (FDR<0.01) for migration and invasion-related processes, including EMT, cell motility and locomotion (**Figure 3A**). Of note, the MM057 melanoma cell line that was used for the RNA-seq experiment harbor a proliferative transcriptome (4). Together, these findings suggest that BORIS contributes to a switch from a proliferative to invasive transcriptome in melanoma cells.

**Figure 3.**
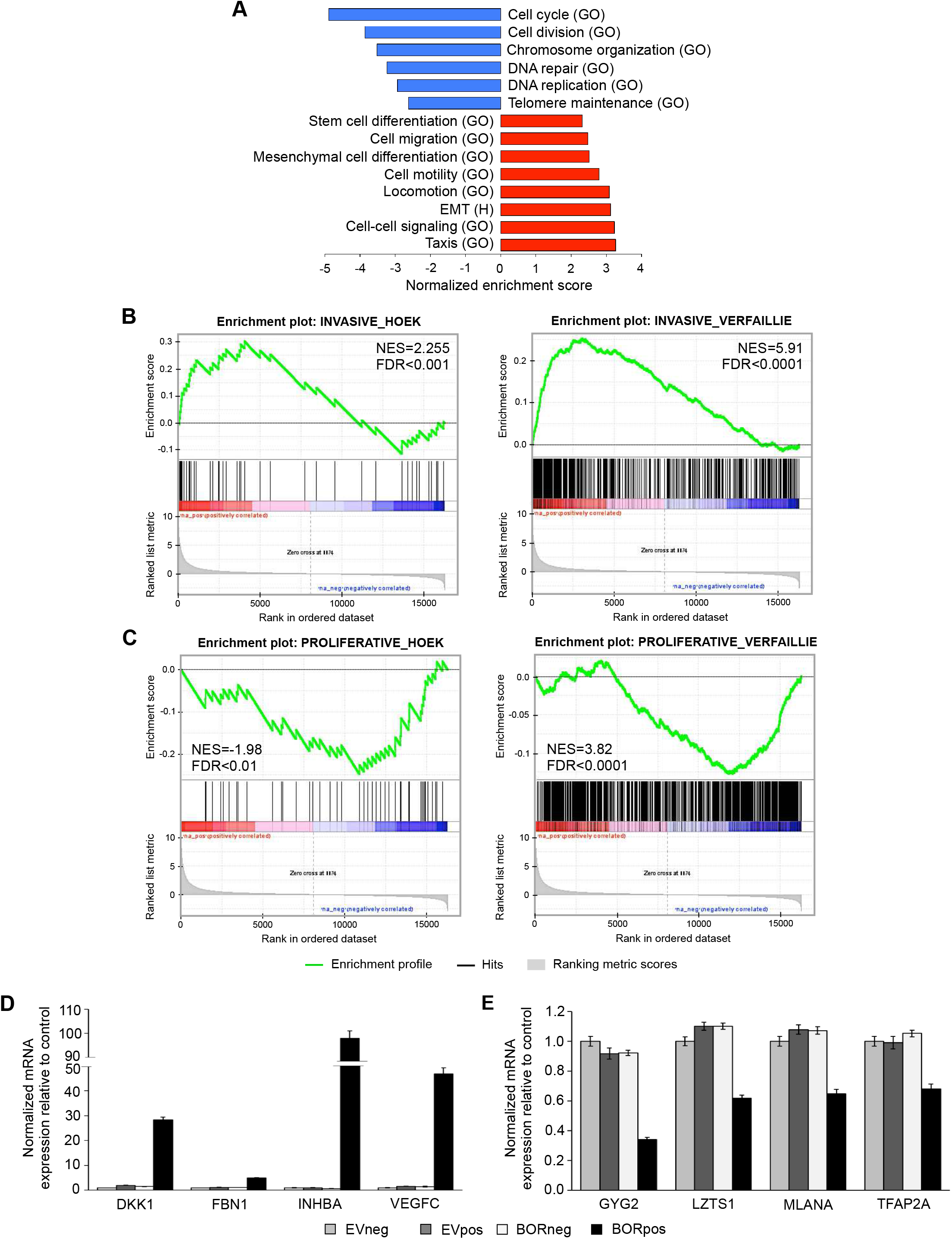
BORIS promotes a switch from a proliferative to invasive gene signature in melanoma cells. **(A-C)** GSEA on the RNA-seq data from the MM057 melanoma cell line for **(A)** biological processes and the hallmark EMT gene set from the Molecular Signatures Database, **(B)** the invasive, and **(C)** proliferative gene signatures. (**D, E)** BORIS expression was induced in the MM057 cell line using 50ng/ml doxycycline for 5 days followed by qPCR for **(D)** invasive and **(E)** proliferative genes. For each sample, a technical triplicate was performed for two biological replicates. Expression was normalized to *HPRT1* and *TBP*. Error bars represent the standard error of mean. GO: gene ontology (biological process), H: Hallmark gene set, NES: normalized enrichment score, FDR: false discovery rate.

To further explore the possibility that BORIS plays a role in phenotype switching we performed GSEA for the melanoma-specific Hoek (3) and Verfaillie (4) gene signatures, which correspond to either the proliferative or invasive transcriptional state of melanoma cells. For these gene sets, we identified a strong positive correlation with the Hoek (NES=2.21, FDR=0.001) and Verfaillie (NES=5.99, FDR<10^−4^) invasive gene signatures (**Figure 3B**) and a negative correlation with the Hoek (NES=−1.98, FDR=0.006) and Verfaillie (NES=−3.84, FDR<10^−4^) proliferative gene signatures (**Figure 3C**). Furthermore, when we compared the upregulated DEGs from the RNA-seq with genes that belong to the invasive and proliferative gene-signatures we observed a significant overlap between invasion-associated genes and the upregulated DEGs (*P*=5.37e-14), but not between the proliferation-associated genes and the upregulated DEGs (*P*=0.28, **Figure S5A**). A similar analysis with the downregulated DEGs revealed a significant overlap with genes that belong to the proliferative signatures (*P*=3.67e-11), but not with genes from the invasive signatures (*P*=0.87, **Figure S5B**). These data indicate that BORIS enables a transcriptional program that is similar to the invasive transcriptional state.

To confirm the effect of BORIS expression on genes within the gene signatures we first induced BORIS expression in the MM057 cell line for five days with 50ng/ml dox. Next, we performed qPCR for four upregulated and four downregulated DEGs that are part of both the Hoek and Verfaillie invasive and proliferative signatures, respectively. Our results demonstrated a significant upregulation of the four selected invasion-associated genes *DKK1, FBN1, INHBA* and *VEGFC* (**Figure 3D**) and a significant downregulation of the four selected proliferation-associated genes *GYG2, LZTS1, MLANA* and *TFAP2A* (**Figure 3E**) in the BORpos cells compared to the control cells. Together, these results indicate that BORIS expression in the MM057 melanoma cell line promotes a switch from a proliferative to an invasive transcriptional state.

### BORIS expression promotes an invasive phenotype

The observation that BORIS can contribute to an invasive transcriptional state prompted us to ask if BORIS expression can promote an invasive phenotype in melanoma cells. To address this question, we first assessed the effect of ectopic BORIS expression on the migration of melanoma cells using a transwell assay. BORIS expression was induced for five days with 100ng/ml dox in two melanoma cell lines (MM057 and MM074) that both harbor a proliferative transcriptional state (4). Compared to the control cells, we observed a significant increase in the percentage of migrating cells upon BORIS expression (**Figure 4A**). Next, we tested the invasive capacity of both cell lines following BORIS expression in a Matrigel transwell invasion assay. Our results demonstrated a significant increase in the invasive abilities of both cell lines in the presence of BORIS (**Figure 4B**). Overall, these results strongly support the idea that BORIS expression leads to an increase in the migratory and invasive potential of proliferative melanoma cell lines.

**Figure 4.**
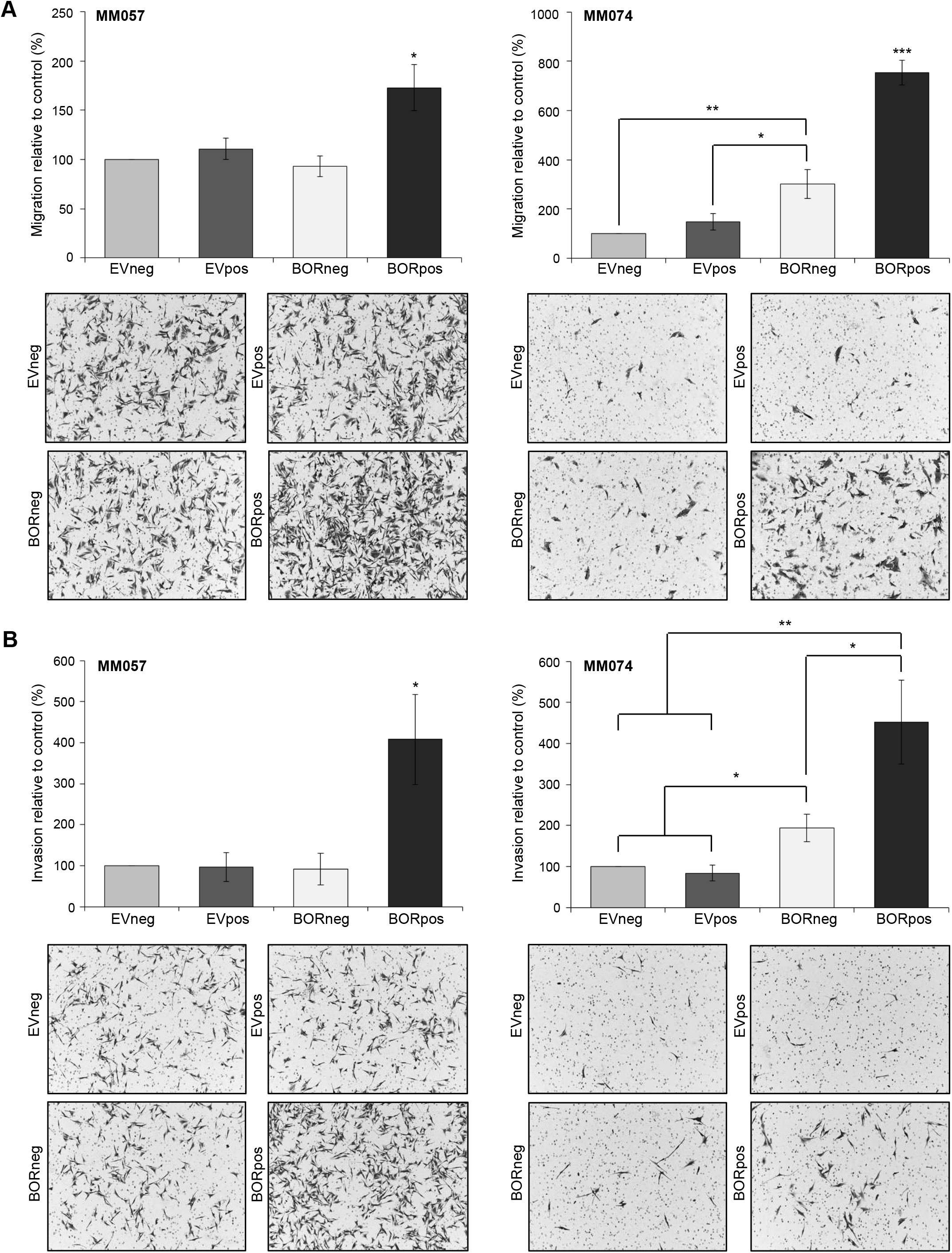
BORIS expression results in increased migration and invasion of melanoma cells with a proliferative gene signature. **(A, B)** BORIS expression was induced in the MM057 and MM074 melanoma cell lines using 100ng/ml doxycycline for 5 days. *In vitro* cell **(A)** migration and **(B)** invasion was determined by transwell assays. Bar graphs display the percentage of cells per field that **(A)** migrated and **(B)** invaded through the membrane and are accompanied by a representative image. For each assay, images of six fields were captured and counted using the Cell Counter option in ImageJ. Three independent experiments were performed, each consisting of two technical replicates. * indicates significantly different from the controls (\**P*<0.05. \*\**P*<0.01, \*\*\**P*<0.001).

### Identification of putative BORIS target genes among invasion-associated genes

To obtain a better understanding of how BORIS might contribute to the observed transcriptional changes, we used *in silico* analyses to identify putative direct BORIS target genes among the DEGs from the RNA-seq data. First, we used iRegulon (39), which identifies master regulators of gene sets based on enrichment for specific DNA binding motifs around the transcription start site. Motif discovery predicted CTCFL (BORIS) as the most enriched transcription factor-binding motif among the upregulated DEGs (NES 3.789). Based on this analysis, 241 putative BORIS target genes were identified, representing 18.4% of all upregulated genes (**Figure 5A**). In contrast, motif discovery did not reveal enrichment for a CTCFL (BORIS) binding motif among downregulated genes. Together, these findings suggest that BORIS may act as a direct transcriptional regulator for a subset of upregulated DEGs.

**Figure 5.**
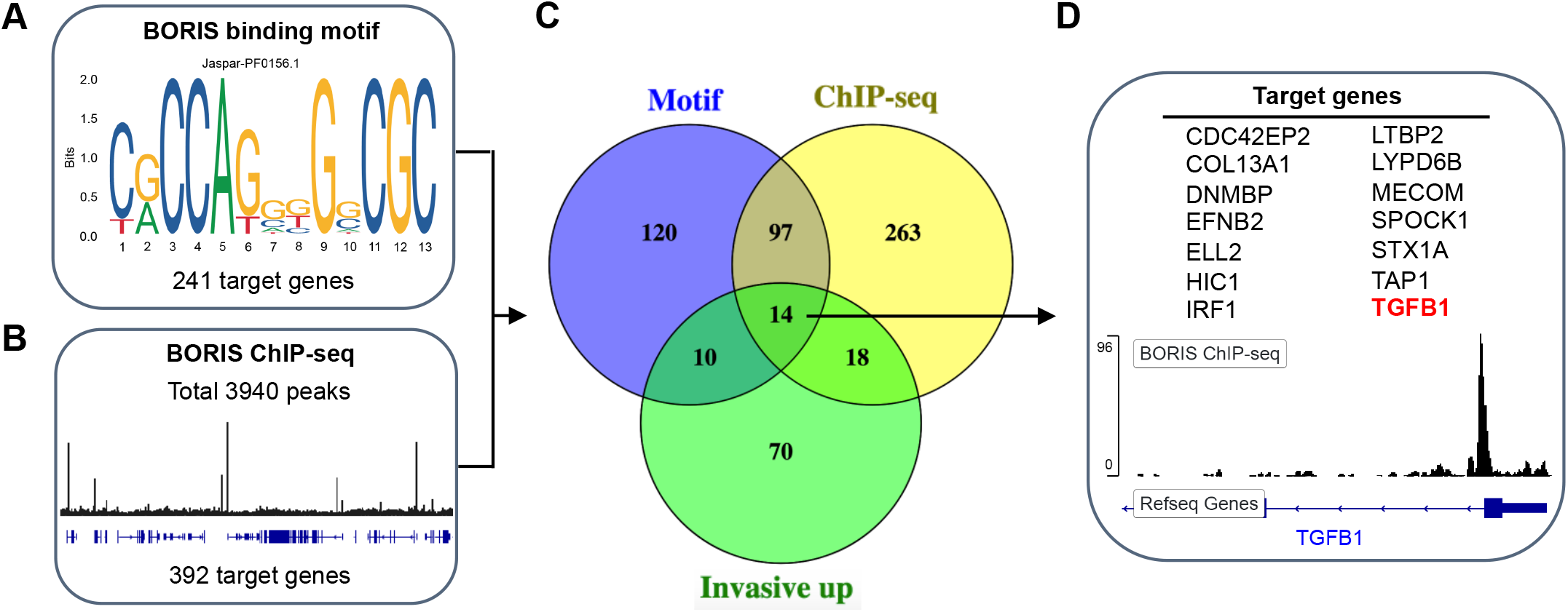
Identification of putative direct BORIS target genes by *in silico* analysis. **(A)** iRegulon was used to assess the presence of the depicted BORIS binding motif around the transcription start site of the DEGs (from RNA-seq). Among upregulated DEGs, 241 putative direct BORIS target genes were identified. **(B)** The BORIS ChIP-seq dataset from ENCODE was provided to the Genomic Regions Enrichment of Annotations Tool, which revealed 3940 BORIS binding sites within 5kb around the transcription start site. The image represents part of the ENCODE BORIS ChIP-seq track as observed within the Integrative Genomics Viewer. Comparing the genes that contain a BORIS binding site with the upregulated DEGs identified 392 putative direct BORIS target genes. **(C)** Venn diagram demonstrating the overlap between putative direct BORIS target genes based on the BORIS binding motif (241 genes), BORIS binding site according to the BORIS-ChIP-seq dataset (392 genes), and upregulated DEGs that belong to the invasive gene signatures (112 genes). **(D)** Overview of the identified 14 putative direct BORIS target genes. The image shows the BORIS binding site in *TGFB1* near the transcription start site based on the ENCODE BORIS ChIP-seq track as observed within the Integrative Genomics Viewer. DEGs: differentially expressed genes, ENCODE: Encyclopedia of DNA Elements.

Next, we used the Genomic Regions Enrichment of Annotations Tool (GREAT) (40) to extend the list of putative direct BORIS target genes. This tool allows the use of cis-regulatory regions obtained from ChIP-seq data to identify nearby annotated genes. As input, we used the publicly available BORIS ChIP-seq dataset from the ENCyclopedia Of DNA Elements (ENCODE) (41). We specifically looked for BORIS binding sites 5kb around the transcription start site. This analysis revealed 3940 genes that were bound in their promoter region by BORIS. From these genes, 392 belonged to the upregulated genes (**Figure 5B**) and 96 to the downregulated genes. Consistent with the results from iRegulon, this analysis revealed a higher number of BORIS binding sites in the gene promoters of the upregulated genes compared to the downregulated genes.

Given that BORIS-DNA binding is more prevalent in the promoter region of genes that become transcriptionally activated (23), we further focused our analysis on the upregulated DEGs. We compared the upregulated putative BORIS target genes as identified via motif discovery (241 genes) and ChIP-seq analysis (392 genes), which revealed 111 genes that contain both a BORIS binding motif and were bound by BORIS in the ChIP-seq dataset (**Figure 5C**). To elucidate putative direct BORIS target genes among genes of the invasive melanoma gene signatures, we compared these genes to the invasion-associated genes that are upregulated by BORIS (112 genes). Based on this analysis we identified 14 potential BORIS target genes that are part of the invasive gene signatures (**Figure 5C**). Interestingly, one of the putative direct BORIS targets in this list is the well-known inducer of melanoma cell phenotype switching (3, 4, 42), TGF-beta (*TGFB1*) (**Figure 5D**). Overall, these analyses suggest that BORIS can directly modulate the expression of a subset of invasion-associated genes, including *TGFB1*.

### BORIS binds the *TGFB1* gene and upregulates *TGFB1* expression

Having identified *TGFB1* as a putative direct BORIS target gene, we set out to validate the *in silico* analysis in melanoma cells. To this end, BORIS binding in the *TGFB1* gene was assessed using chromatin immunoprecipitation (ChIP) followed by qPCR. First we generated a dox-inducible expression construct coding for BORIS fused to a triple FLAG-tag, as commercially available BORIS antibodies are not suitable for ChIP. BORIS expression was induced for five days with 100ng/ml dox in two melanoma cell lines (MM057 and MM074) and expression was confirmed by both immunoblot (**Figure 6A**). Next, primers were designed for ChIP-qPCR based on the consensus DNA sequence surrounding the putative BORIS binding site near the *TGFB1* promoter (**Figure 5D**). In addition, primers recognizing an intergenic region depleted of putative BORIS/CTCF binding sites were designed and served as negative control. Compared to the controls, we observed a significant enrichment in the immunoprecipitation of BORIS in the *TGFB1* gene in both melanoma cell lines (**Figure 6B**), demonstrating that BORIS physically binds the identified binding site in close proximity of the *TGFB1* promoter.

**Figure 6.**
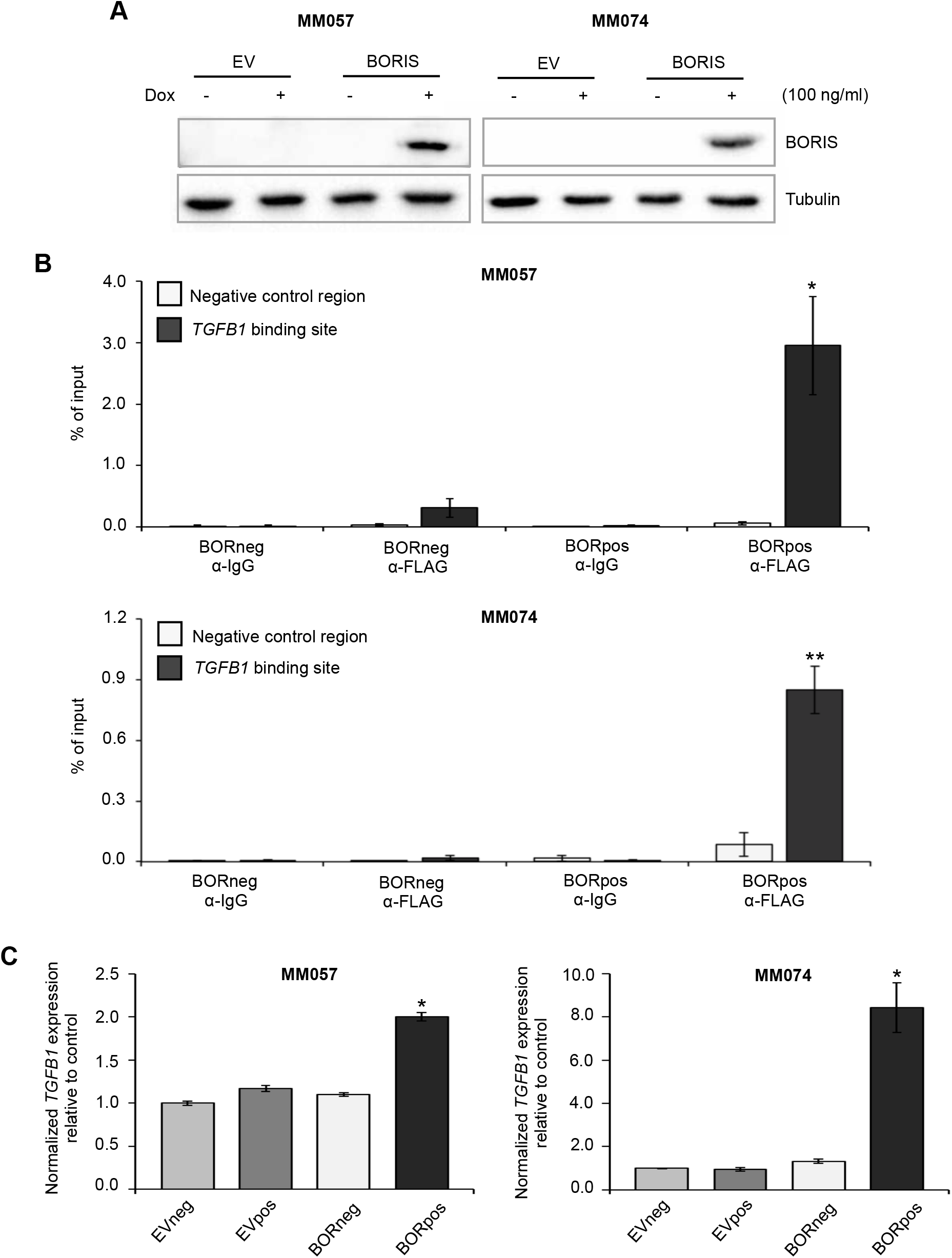
BORIS binds near the *TGFB1* promoter and upregulates *TGFB1* expression in melanoma cells. **(A-C)** Expression of BORIS fused to a triple FLAG-tag was induced in the MM057 and MM074 melanoma cell lines with 100ng/ml dox for 5 days. **(A)** Whole cell lysate was used for immunoblotting with anti-BORIS antibody. Anti-Tubulin was used as a loading control. **(B)** ChIP with either anti-FLAG antibody or mouse IgG followed by qPCR for either the putative BORIS binding site in *TGFB1* or a negative control region ∼3kbp upstream of the putative binding site. **(C)** qPCR for *TGFB1*. For each qPCR experiment, a technical triplicate was performed for two biological replicates. Expression was normalized to *HPRT1* and *TBP*. Error bars represent the standard error of the mean. * indicates significantly different from controls (\**P*<0.05. \*\**P*<0.01). Dox: doxycycline

Next, we wanted to assess if BORIS-DNA binding in *TGFB1* is accompanied by a change in *TGFB1* expression. Using qPCR, we found increased *TGFB1* levels upon BORIS expression compared to the controls in both the MM057 and MM074 cell lines (**Figure 6C**), which is consistent with the RNA-seq data. These findings suggest that BORIS binding near the *TGFB1* promoter leads to upregulation of *TGFB1* expression. In addition to *TGFB1*, our RNA-seq data revealed an increase in the expression of various genes belonging to the TGF-beta ligand family, including activins, bone morphogenetic proteins, and growth and differentiation factors (**Figure S6A**). Furthermore, multiple TGF-beta target genes are significantly upregulated in the RNA-seq dataset, including *FBN1*, *SERPINE1*, *JUNB*, *FOSB*, and *NGFR/CD271* (**Figure S6B**). In addition, *SNAI1* is upregulated in the RNA-seq dataset, though not significantly (FDR=0.08, **Figure S6B**) Importantly, *CD271*, *SNAI1*, *JUNB* and *FOSB*, the latter two as part of the AP-1 transcription factor complex, are all known regulators of melanoma phenotype switching (4, 8, 43). Together, these data support a direct role for BORIS in the expression of the EMT-inducer TGF-beta and TGF-beta target genes, revealing one mechanism through which BORIS could modulate the transcription state and contribute to an invasive phenotype in melanoma cells.

## Discussion

BORIS is a DNA-binding protein with high similarity to the multifunctional transcription factor CTCF. In contrast to ubiquitously expressed CTCF, BORIS expression is restricted to the testis (12) and becomes aberrantly re-activated in various types of cancer, including melanoma (34). Currently, the role of BORIS in melanoma remains elusive. In this study, we demonstrate that BORIS can alter the gene expression program of melanoma cells in favor of a more invasive phenotype. Transcriptional profiling revealed that genes upregulated in melanoma cells with ectopic BORIS expression are enriched among established invasive gene signatures, while downregulated genes are enriched among the proliferative gene signatures. In line with these findings, we showed that BORIS expression resulted in reduced proliferation, whereas the migratory and invasive abilities of melanoma cells were enhanced. Furthermore, the upregulation of various TGF-beta family members and TGF-beta target genes in BORIS-expressing cells together with BORIS’ ability to bind in close proximity to the *TGFB1* promoter in melanoma cells, indicates a direct role for BORIS in melanoma phenotype switching via TGF-beta. Together, our data indicate that expression of BORIS contributes to the switch from a proliferative towards an invasive state at both the transcriptional and phenotypic level in melanoma cells.

Recent studies reported BORIS expression in female germ cells and various somatic tissues (34, 44), which challenges the initial description of BORIS as a cancer testis antigen. If BORIS is indeed expressed at the protein level in somatic tissues remains a debate (24). At the RNA level, the GTEx dataset demonstrates *BORIS* expression in most normal tissues, though at a much lower level compared to testis. In normal skin we found that *BORIS* expression is very low with detectable levels of BORIS in only one third of the samples, while expression in melanoma is significantly higher. Along the same lines, we were unable to detect *BORIS* expression in three, non-malignant giant nevi cell lines, while we did measure *BORIS* expression in most of our melanoma cell lines. These observations are in agreement with a previous study where expression of BORIS was measured in melanoma cell lines and samples, but not in normal skin tissue (36). Unfortunately, due to the lack of a good commercially available BORIS antibody we were unable to reliably detect endogenous BORIS protein by either immunoblot or immunohistochemistry in melanoma cell lines and tissue.

BORIS has previously been shown to be involved in numerous cellular processes (34, 45), many of which are altered during carcinogenesis. Here, we found that BORIS expression leads to decreased proliferation and increased apoptosis. The decrease in proliferation that we observed is in agreement with previous studies that report reduced proliferation upon ectopic BORIS expression in both primary cells and various cancer cell lines (46, 47). Though the literature regarding proliferation in the context of BORIS is not uniform, as others have reported increased proliferation upon BORIS overexpression (17, 20, 37). Similarly, previous studies have revealed different effects of altered BORIS expression on apoptosis (16, 37, 46, 48). These observations indicate a cell type specific function of BORIS, which may be mediated by differences in genome-wide BORIS-DNA binding in different cell types as recently suggested (23).

In this study, we used RNA-seq to gain insight into cellular processes that BORIS might be involved in and that contribute to melanoma development and progression. Our finding that altered BORIS expression leads to large-scale changes in gene expression was expected, since BORIS is known to act as a transcriptional regulator (17, 18, 22, 26). To verify that the transcriptional changes observed upon altered BORIS expression are not a result from BORIS-induced changes in CTCF level, as observed in normal human dermal fibroblasts and ovarian cancer cells (19, 49), we verified a lack of association between BORIS and CTCF expression in both melanoma cell lines and melanoma samples. Interestingly, previous studies reported co-expression between BORIS and various cancer antigens, including members of the melanoma antigen family (28, 29). Our RNA-seq data revealed no association between BORIS and the expression of members of the melanoma antigen family, which is similar to findings in melanoma tissue samples (36). This further supports a cell type specific role for BORIS in cancer.

The gene expression changes observed upon altered BORIS expression indicated a role for BORIS in transcriptional reprogramming of melanoma cells from a proliferative to an invasive state. This reversible process, known as phenotype switching, is critical for melanoma cells to acquire an invasive phenotype that fuels disease progression (50). In line with the acquisition of a pro-invasive transcriptional state, we demonstrated that BORIS-expressing melanoma cells display reduced proliferation and increased migratory and invasive abilities. Involvement of BORIS in EMT, a process in epithelial cells that is similar to phenotype switching, was observed in liver cancer cell lines (18), human fallopian tube secretory epithelial cells (51) as well as the CD44/CD133 positive subpopulation of the IMR-32 neuroblastoma cell line (52). However, in contrast, one study showed that BORIS knockdown promotes EMT in MCF7 breast cancer cells (53). Multiple studies found that increased BORIS expression is linked to poor prognosis and more advanced stages of disease, hence BORIS being considered an oncogene (54–57). Our data indicates a pro-invasive role for BORIS in melanoma and supports the notion that BORIS acts pro-oncogenic.

The large number of genes that belong to the proliferative and invasive gene signatures highlights the complexity that underlies the process of phenotype switching. Multiple transcription factors, signalling pathways and EMT-related genes are important to establish the transitions (3, 4, 8–11). Trying to understand which factors contribute to the BORIS-induced transcriptional switch towards an invasive state, we used two complementary *in silico* approaches to identified putative direct BORIS target genes. Of note, iRegulon, which was used to identify targets based on the BORIS binding motif, has been used previously to identify regulators of phenotype switching (4). Since the only ENCODE ChIP-seq dataset for BORIS is derived from a myeloid leukemia cell line (K562), we cannot exclude the possibility that the analysis missed certain melanoma-specific direct BORIS target genes. Nevertheless, this analysis did reveal a larger number of potential direct BORIS target genes compared to the DNA binding motif-based analysis. In agreement with the observation that BORIS-DNA binding is enriched at chromatin regions marked by activating histone modifications (23), the majority of the identified putative direct BORIS target genes were among the upregulated DEGs.

In the current study, we focused our validation experiments of the *in silico* determined direct BORIS target genes on *TGFB1*, since TGF-beta driven signaling is an important mark of the invasive signature (1, 4), TGF-beta acts as tumor promoter at later stages of tumor development (58) and CTCF was predicted to bind within the *TGFB1* regulatory region and function as a transcriptional regulator of *TGFB1* expression (59). Here, we demonstrate the ability of BORIS to bind in the vicinity of the *TGFB1* promoter and show that ectopic BORIS expression leads to increased expression of *TGFB1*. In addition, the expression of TGF-beta family members and TGF-beta target genes was increased, which is likely mediated by autocrine and paracrine signaling between tumor cells. Interestingly, a previous study that characterized *Ctcfl* transgenic mice observed a phenotype that resembled mice with an altered TGF-beta pathway. In addition, RNA-seq of *Ctcfl* transgenic embryonic stem cells revealed disruption of the TGF-beta pathway as well as increased *TgfB1* expression (60), which is in line with our observations. Importantly, among the TGF-beta target genes that were upregulated in our RNA-seq dataset are *CD271*, *SNAI1*, and *JUNB*, which are all known regulators of melanoma phenotype switching (4, 8, 43). These observations suggest that BORIS acts as a transcriptional regulator of phenotype switching via *TGFB1* and warrant further investigations into the role of BORIS in TGF-beta signaling.

Based on our findings, we propose that BORIS expression induces phenotype switching via upregulation of TGF-beta signaling. In addition to the regulation of *TGFB1* by BORIS, other mechanisms might contribute to BORIS-mediated transcriptional changes, including competition with CTCF for DNA binding sites, epigenetic modifications, and/or chromatin looping, which are the subject of further investigations. Taken together, our study demonstrates that BORIS contributes to melanoma progression by intrinsically rewiring gene expression to promote an invasive phenotype. Furthermore, these results indicate that BORIS is an important transcriptional regulator during phenotype switching in melanoma and provide a rationale for further studies into BORIS’ role as an invasion promoting transcriptional regulator.

## Acknowledgements

This work was partially funded by the Israel Cancer Research Fund. SMJ was supported by the Banque National Fellowship, the Dr. Victor K.S. Lui Fellowship and the CIHR/FRSQ training grant in cancer research (FRN53888) of the McGill Integrated Cancer Research Training Program. AIP was supported by the Banque National Fellowship.

## References

1. Hoek KS, Eichhoff OM, Schlegel NC, Dobbeling U, Kobert N, Schaerer L, et al. In vivo switching of human melanoma cells between proliferative and invasive states. Cancer Res. 2008;68(3):650–6.

2. Hoek KS, Goding CR. Cancer stem cells versus phenotype-switching in melanoma. Pigment Cell Melanoma Res. 2010;23(6):746–59.

3. Hoek KS, Schlegel NC, Brafford P, Sucker A, Ugurel S, Kumar R, et al. Metastatic potential of melanomas defined by specific gene expression profiles with no BRAF signature. Pigment Cell Res. 2006;19(4):290–302.

4. Verfaillie A, Imrichova H, Atak ZK, Dewaele M, Rambow F, Hulselmans G, et al. Decoding the regulatory landscape of melanoma reveals TEADS as regulators of the invasive cell state. Nat Commun. 2015;6:6683.

5. Widmer DS, Cheng PF, Eichhoff OM, Belloni BC, Zipser MC, Schlegel NC, et al. Systematic classification of melanoma cells by phenotype-specific gene expression mapping. Pigment Cell Melanoma Res. 2012;25(3):343–53.

6. Jeffs AR, Glover AC, Slobbe LJ, Wang L, He S, Hazlett JA, et al. A gene expression signature of invasive potential in metastatic melanoma cells. PLoS One. 2009;4(12):e8461.

7. Tirosh I, Izar B, Prakadan SM, Wadsworth MH, 2nd, Treacy D, Trombetta JJ, et al. Dissecting the multicellular ecosystem of metastatic melanoma by single-cell RNA-seq. Science (New York, NY). 2016;352(6282):189–96.

8. Aibar S, Gonzalez-Blas CB, Moerman T, Huynh-Thu VA, Imrichova H, Hulselmans G, et al. SCENIC: single-cell regulatory network inference and clustering. Nat Methods. 2017;14(11):1083–6.

9. Fane ME, Chhabra Y, Smith AG, Sturm RA. BRN2, a POUerful driver of melanoma phenotype switching and metastasis. Pigment Cell Melanoma Res. 2018.

10. Pinner S, Jordan P, Sharrock K, Bazley L, Collinson L, Marais R, et al. Intravital imaging reveals transient changes in pigment production and Brn2 expression during metastatic melanoma dissemination. Cancer Res. 2009;69(20):7969–77.

11. Perotti V, Baldassari P, Molla A, Nicolini G, Bersani I, Grazia G, et al. An actionable axis linking NFATc2 to EZH2 controls the EMT-like program of melanoma cells. Oncogene. 2019.

12. Loukinov DI, Pugacheva E, Vatolin S, Pack SD, Moon H, Chernukhin I, et al. BORIS, a novel male germ-line-specific protein associated with epigenetic reprogramming events, shares the same 11-zinc-finger domain with CTCF, the insulator protein involved in reading imprinting marks in the soma. Proc Natl Acad Sci U S A. 2002;99(10):6806–11.

13. Phillips JE, Corces VG. CTCF: master weaver of the genome. Cell. 2009;137(7):1194–211.

14. Kemp CJ, Moore JM, Moser R, Bernard B, Teater M, Smith LE, et al. CTCF haploinsufficiency destabilizes DNA methylation and predisposes to cancer. Cell Rep. 2014;7(4):1020–9.

15. Rasko JE, Klenova EM, Leon J, Filippova GN, Loukinov DI, Vatolin S, et al. Cell growth inhibition by the multifunctional multivalent zinc-finger factor CTCF. Cancer Res. 2001;61(16):6002–7.

16. Dougherty CJ, Ichim TE, Liu L, Reznik G, Min WP, Ghochikyan A, et al. Selective apoptosis of breast cancer cells by siRNA targeting of BORIS. Biochem Biophys Res Commun. 2008;370(1):109–12.

17. Gaykalova D, Vatapalli R, Glazer CA, Bhan S, Shao C, Sidransky D, et al. Dose-dependent activation of putative oncogene SBSN by BORIS. PLoS One. 2012;7(7):e40389.

18. Liu Q, Chen K, Liu Z, Huang Y, Zhao R, Wei L, et al. BORIS up-regulates OCT4 via histone methylation to promote cancer stem cell-like properties in human liver cancer cells. Cancer Lett. 2017;403:165–74.

19. Renaud S, Loukinov D, Alberti L, Vostrov A, Kwon YW, Bosman FT, et al. BORIS/CTCFL-mediated transcriptional regulation of the hTERT telomerase gene in testicular and ovarian tumor cells. Nucleic Acids Res. 2011;39(3):862–73.

20. Smith IM, Glazer CA, Mithani SK, Ochs MF, Sun W, Bhan S, et al. Coordinated activation of candidate proto-oncogenes and cancer testes antigens via promoter demethylation in head and neck cancer and lung cancer. PLoS One. 2009;4(3):e4961.

21. Sun L, Huang L, Nguyen P, Bisht KS, Bar-Sela G, Ho AS, et al. DNA methyltransferase 1 and 3B activate BAG-1 expression via recruitment of CTCFL/BORIS and modulation of promoter histone methylation. Cancer Res. 2008;68(8):2726–35.

22. Zampieri M, Ciccarone F, Palermo R, Cialfi S, Passananti C, Chiaretti S, et al. The epigenetic factor BORIS/CTCFL regulates the NOTCH3 gene expression in cancer cells. Biochim Biophys Acta. 2014;1839(9):813–25.

23. Bergmaier P, Weth O, Dienstbach S, Boettger T, Galjart N, Mernberger M, et al. Choice of binding sites for CTCFL compared to CTCF is driven by chromatin and by sequence preference. Nucleic Acids Res. 2018;46(14):7097–107.

24. Pugacheva EM, Rivero-Hinojosa S, Espinoza CA, Mendez-Catala CF, Kang S, Suzuki T, et al. Comparative analyses of CTCF and BORIS occupancies uncover two distinct classes of CTCF binding genomic regions. Genome Biol. 2015;16:161.

25. Sleutels F, Soochit W, Bartkuhn M, Heath H, Dienstbach S, Bergmaier P, et al. The male germ cell gene regulator CTCFL is functionally different from CTCF and binds CTCF-like consensus sites in a nucleosome composition-dependent manner. Epigenetics Chromatin. 2012;5(1):8.

26. Hong JA, Kang Y, Abdullaev Z, Flanagan PT, Pack SD, Fischette MR, et al. Reciprocal binding of CTCF and BORIS to the NY-ESO-1 promoter coincides with derepression of this cancer-testis gene in lung cancer cells. Cancer Res. 2005;65(17):7763–74.

27. Kang Y, Hong JA, Chen GA, Nguyen DM, Schrump DS. Dynamic transcriptional regulatory complexes including BORIS, CTCF and Sp1 modulate NY-ESO-1 expression in lung cancer cells. Oncogene. 2007;26(30):4394–403.

28. Bhan S, Negi SS, Shao C, Glazer CA, Chuang A, Gaykalova DA, et al. BORIS binding to the promoters of cancer testis antigens, MAGEA2, MAGEA3, and MAGEA4, is associated with their transcriptional activation in lung cancer. Clin Cancer Res. 2011;17(13):4267–76.

29. Vatolin S, Abdullaev Z, Pack SD, Flanagan PT, Custer M, Loukinov DI, et al. Conditional expression of the CTCF-paralogous transcriptional factor BORIS in normal cells results in demethylation and derepression of MAGE-A1 and reactivation of other cancer-testis genes. Cancer Res. 2005;65(17):7751–62.

30. Jelinic P, Stehle JC, Shaw P. The testis-specific factor CTCFL cooperates with the protein methyltransferase PRMT7 in H19 imprinting control region methylation. PLoS Biol. 2006;4(11):e355.

31. Nguyen P, Bar-Sela G, Sun L, Bisht KS, Cui H, Kohn E, et al. BAT3 and SET1A form a complex with CTCFL/BORIS to modulate H3K4 histone dimethylation and gene expression. Mol Cell Biol. 2008;28(21):6720–9.

32. Lobanenkov VV, Zentner GE. Discovering a binary CTCF code with a little help from BORIS. Nucleus. 2018;9(1):33–41.

33. Martin-Kleiner I. BORIS in human cancers -- a review. Eur J Cancer. 2012;48(6):929–35.

34. Soltanian S, Dehghani H. BORIS: a key regulator of cancer stemness. Cancer Cell Int. 2018;18:154.

35. Klenova EM, Morse HC, Ohlsson R, Lobanenkov VV. The novel BORIS + CTCF gene family is uniquely involved in the epigenetics of normal biology and cancer. Seminars in Cancer Biology. 2002;12(5):399–414.

36. Kholmanskikh O, Loriot A, Brasseur F, De Plaen E, De Smet C. Expression of BORIS in melanoma: lack of association with MAGE-A1 activation. Int J Cancer. 2008;122(4):777–84.

37. Zhang Y, Fang M, Song Y, Ren J, Fang J, Wang X. Brother of Regulator of Imprinted Sites (BORIS) suppresses apoptosis in colorectal cancer. Sci Rep. 2017;7:40786.

38. Subramanian A, Tamayo P, Mootha VK, Mukherjee S, Ebert BL, Gillette MA, et al. Gene set enrichment analysis: a knowledge-based approach for interpreting genome-wide expression profiles. Proc Natl Acad Sci U S A. 2005;102(43):15545–50.

39. Janky R, Verfaillie A, Imrichova H, Van de Sande B, Standaert L, Christiaens V, et al. iRegulon: from a gene list to a gene regulatory network using large motif and track collections. PLoS computational biology. 2014;10(7):e1003731.

40. McLean CY, Bristor D, Hiller M, Clarke SL, Schaar BT, Lowe CB, et al. GREAT improves functional interpretation of cis-regulatory regions. Nat Biotechnol. 2010;28(5):495–501.

41. Davis CA, Hitz BC, Sloan CA, Chan ET, Davidson JM, Gabdank I, et al. The Encyclopedia of DNA elements (ENCODE): data portal update. Nucleic Acids Res. 2018;46(D1):D794–d801.

42. Li FZ, Dhillon AS, Anderson RL, McArthur G, Ferrao PT. Phenotype switching in melanoma: implications for progression and therapy. Front Oncol. 2015;5:31.

43. Restivo G, Diener J, Cheng PF, Kiowski G, Bonalli M, Biedermann T, et al. low neurotrophin receptor CD271 regulates phenotype switching in melanoma. Nat Commun. 2017;8(1):1988.

44. Jones TA, Ogunkolade BW, Szary J, Aarum J, Mumin MA, Patel S, et al. Widespread expression of BORIS/CTCFL in normal and cancer cells. PLoS One. 2011;6(7):e22399.

45. Marshall AD, Bailey CG, Rasko JE. CTCF and BORIS in genome regulation and cancer. Curr Opin Genet Dev. 2014;24:8–15.

46. Tiffen JC, Bailey CG, Marshall AD, Metierre C, Feng Y, Wang Q, et al. The cancer-testis antigen BORIS phenocopies the tumor suppressor CTCF in normal and neoplastic cells. Int J Cancer. 2013;133(7):1603–13.

47. Rosa-Garrido M, Ceballos L, Alonso-Lecue P, Abraira C, Delgado MD, Gandarillas A. A cell cycle role for the epigenetic factor CTCF-L/BORIS. PLoS One. 2012;7(6):e39371.

48. Alberti L, Renaud S, Losi L, Leyvraz S, Benhattar J. High expression of hTERT and stemness genes in BORIS/CTCFL positive cells isolated from embryonic cancer cells. PLoS One. 2014;9(10):e109921.

49. Renaud S, Pugacheva EM, Delgado MD, Braunschweig R, Abdullaev Z, Loukinov D, et al. Expression of the CTCF-paralogous cancer-testis gene, brother of the regulator of imprinted sites (BORIS), is regulated by three alternative promoters modulated by CpG methylation and by CTCF and p53 transcription factors. Nucleic Acids Res. 2007;35(21):7372–88.

50. Caramel J, Papadogeorgakis E, Hill L, Browne GJ, Richard G, Wierinckx A, et al. A switch in the expression of embryonic EMT-inducers drives the development of malignant melanoma. Cancer Cell. 2013;24(4):466–80.

51. Hillman JC, Pugacheva EM, Barger CJ, Sribenja S, Rosario S, Albahrani M, et al. BORIS expression in ovarian cancer precursor cells alters the CTCF cistrome and enhances invasiveness through GALNT14. Mol Cancer Res. 2019.

52. Garikapati KR, Patel N, Makani VKK, Cilamkoti P, Bhadra U, Bhadra MP. Down-regulation of BORIS/CTCFL efficiently regulates cancer stemness and metastasis in MYCN amplified neuroblastoma cell line by modulating Wnt/beta-catenin signaling pathway. Biochem Biophys Res Commun. 2017;484(1):93–9.

53. Alberti L, Losi L, Leyvraz S, Benhattar J. Different Effects of BORIS/CTCFL on Stemness Gene Expression, Sphere Formation and Cell Survival in Epithelial Cancer Stem Cells. PLoS One. 2015;10(7):e0132977.

54. Cheema Z, Hari-Gupta Y, Kita GX, Farrar D, Seddon I, Corr J, et al. Expression of the cancer-testis antigen BORIS correlates with prostate cancer. Prostate. 2014;74(2):164–76.

55. Chen K, Huang W, Huang B, Wei Y, Li B, Ge Y, et al. BORIS, brother of the regulator of imprinted sites, is aberrantly expressed in hepatocellular carcinoma. Genet Test Mol Biomarkers. 2013;17(2):160–5.

56. D’Arcy V, Abdullaev ZK, Pore N, Docquier F, Torrano V, Chernukhin I, et al. The potential of BORIS detected in the leukocytes of breast cancer patients as an early marker of tumorigenesis. Clin Cancer Res. 2006;12(20 Pt 1):5978–86.

57. Woloszynska-Read A, Zhang W, Yu J, Link PA, Mhawech-Fauceglia P, Collamat G, et al. Coordinated cancer germline antigen promoter and global DNA hypomethylation in ovarian cancer: association with the BORIS/CTCF expression ratio and advanced stage. Clin Cancer Res. 2011;17(8):2170–80.

58. Perrot CY, Javelaud D, Mauviel A. Insights into the Transforming Growth Factor-beta Signaling Pathway in Cutaneous Melanoma. Annals of dermatology. 2013;25(2):135–44.

59. Dhaouadi N, Li JY, Feugier P, Gustin MP, Dab H, Kacem K, et al. Computational identification of potential transcriptional regulators of TGF-ss1 in human atherosclerotic arteries. Genomics. 2014;103(5-6):357–70.

60. Sati L, Zeiss C, Yekkala K, Demir R, McGrath J. Expression of the CTCFL Gene during Mouse Embryogenesis Causes Growth Retardation, Postnatal Lethality, and Dysregulation of the Transforming Growth Factor beta Pathway. Mol Cell Biol. 2015;35(19):3436–45.

